# Chronic Cochlear Implantation with and without Electric Stimulation in a Mouse Model Induces Robust Cochlear Influx of CX3CR1^+/GFP^ Macrophages

**DOI:** 10.1101/2021.10.31.466540

**Authors:** Alexander D. Claussen, René Vielman Quevedo, Timon Higgins, Brian Mostaert, Muhammad Taifur Rahman, Jonathon Kirk, Keiko Hirose, Marlan R. Hansen

## Abstract

**Background:** Cochlear implantation is an effective auditory rehabilitation strategy for those with profound hearing loss, including those with residual low frequency hearing through use of hybrid cochlear implantation techniques. Post-mortem studies demonstrate the nearly ubiquitous presence of intracochlear fibrosis and neo-ossification following cochlear implantation. Current evidence suggests post-implantation intracochlear fibrosis is associated with delayed loss of residual acoustic hearing in hybrid cochlear implant (CI) recipients and may also negatively influence outcomes in traditional CI recipients. This study examined the contributions of surgical trauma, foreign body response and electric stimulation to intracochlear fibrosis and the innate immune response to cochlear implantation and the hierarchy of these contributions.

**Methods:** Normal hearing CX3CR1^+/GFP^ mice underwent either round window opening (sham), acute CI insertion or chronic CI insertion with no, low- or high-level electric stimulation. Electric stimulation levels were based on neural response telemetry (NRT), beginning post-operative day 7 for 4 hours per day. Subjects (n=3 per timepoint) were sacrificed at 4 hours, 1,4,7,8,11,14 and 21 days. An unimplanted group (n=3) served as controls. Cochleae were harvested at each time-point and prepared for immunohistochemistry with confocal imaging. The images were analyzed to obtain CX3CR1+ macrophage cell number and density in the lateral wall (LW), scala tympani (ST) and Rosenthal’s canal (RC).

**Results:** A ST peri-implant cellular infiltrate and fibrosis occurred exclusively in the chronically implanted groups starting on day 7 with a concurrent infiltration of CX3CR1+ macrophages not seen in the other groups. CX3CR1+ macrophage infiltration was seen in the LW and RC in all experimental groups within the first week, being most prominent in the 3 chronically implanted groups during the second and third week. There were no significant differences in macrophage infiltration related to levels of electric stimulation.

**Conclusions:** The cochlear immune response was most prominent in the presence of chronic cochlear implantation, regardless of electric stimulation level. Further, the development of intracochlear ST fibrosis was dependent on the presence of the indwelling CI foreign body. An innate immune response was evoked by surgical trauma alone (sham and acute CI groups) to a lesser degree. These data suggest that cochlear inflammation and intrascalar fibrosis after cochlear implantation are largely dependent on the presence of a chronic indwelling foreign body and are not critically dependent on electrical stimulation. Also, these data support a role for surgical trauma in inciting the initial innate immune response.

## 1 Introduction

Conventional and hybrid cochlear implantation is an effective treatment for patients with severe and profound hearing loss, including those with preserved low frequency hearing. Advances in cochlear implant (CI) design, surgical technique and programming strategies have improved the hearing performance in CI recipients (Mitchell-Innes et al., 2018; Roche & Hansen, 2015). However, several issues hampering CI efficacy remain, including the development of a post-implantation intracochlear tissue response which may contribute to poorer outcomes in both traditional and hybrid CI recipients (Kamakura & Nadol, 2016; Quesnel et al., 2016; Scheperle et al., 2017). The clinical significance of addressing these issues is increased when considering the anticipated near doubling of the combined conventional and hybrid CI candidate population in the next 40 years for those age 60 years and older (Goman et al., 2018).

Fibrosis and neo-ossification with inflammatory cell infiltration in the human cochlea has been well described in post-implantation cadaveric temporal bone studies, occurring in up to 96% of specimens in some series (Foggia et al., 2019). This heterotopic tissue response is often most robust in the peri-implant region of the scala tympani (ST) forming a ‘fibrous sheath’ around the CI, but is sometimes seen to extend distal to the implant tip and into other scala (Linthicum et al., 2017; Nadol et al., 2014; Seyyedi & Nadol, 2014). A similar pattern of post-implantation cochlear tissue response has been seen in several animal models of cochlear implantation, including guinea pig, cat, mouse and sheep (Clark et al., 1975; Claussen et al., 2019; Kaufmann et al., 2020; O’Leary et al., 2013). Deleterious CI outcomes have been associated with the cochlear tissue response and neo-ossification, including poorer word recognition scores (Kamakura & Nadol, 2016), impedance increases with subsequent poorer battery life and decreased dynamic range (Needham et al., 2020; Wilk et al., 2016) and loss of residual acoustic hearing after cochlear implantation (Quesnel et al., 2016; Scheperle et al., 2017).

A wide variety of inflammatory cells, including lymphocytes, macrophages, eosinophils and peri-implant foreign body giant cells are consistently seen within the intracochlear tissue response as well as other parts of the cochlea (O’Malley et al., 2017). Recent observations from cadaveric temporal bones of previously implanted subjects have demonstrated the presence of macrophages of varying morphologies within the scalae as well as other regions of the cochlea, including Rosenthal’s canal (RC), the osseus spiral lamina and vestibular epithelium (Okayasu et al., 2019, 2020). These observations are limited to the longer post-implantation time-points seen in cadaveric temporal bones, but they demonstrate the presence of a persistent cochlear inflammatory infiltrate following cochlear implantation that may contribute to the formation of the intracochlear tissue response. The initial innate immune and inflammatory response to cochlear implantation is still poorly understood as are the relative contributions of the initial inciting events, including insertional trauma, autologous tissue packing, electric stimulation, foreign body response (FBR) to CI materials and likely other genetic and environmental pre-dispositions (Ishai et al., 2017; O’Leary et al., 2013; Rowe et al., 2016). Understanding of this initial inflammatory response and contributing factors will be important in the development of strategies to mitigate the intracochlear tissue response and improve CI efficacy.

This study utilizes our previously published mouse model of chronic cochlear implantation with electric stimulation in a CX3CR1^+/GFP^ reporter mouse to study the initial innate immune response to cochlear implantation (Claussen et al., 2019). CX3CR1 is the receptor to the chemokine, fractalkine (CX3CL1) and is expressed in monocytes, macrophages, microglia, NK cells and some T cells. (Jung et al., 2000). In the mouse cochlea, CX3CR1+ cells are observed routinely and comprise a population of highly inducible resident cochlear macrophages (Hirose et al., 2005; Sato et al., 2008). Prior studies in CX3CR1^+/GFP^ mice, in which one copy of the *CX3CR1* gene is replaced with a GFP reporter gene, have shown cochlear migration of CX3CR1+ macrophages in response to ototoxic and hair cell injury (Hirose et al., 2014). Further, CX3CR1 knockout studies have demonstrated a protective role of CX3CR1+ monocytes and macrophages following cochlear insult (Kaur et al., 2015; Sato et al., 2010).

Our main objective was to observe the initial, innate immune response to cochlear implantation at several timepoints by tracking the accumulation of CX3CR1+ macrophages within the cochlea. Additionally, we evaluated the relative contributions of several factors to the resultant cochlear immune response, including round window opening, insertional trauma, chronic placement of the CI foreign body and varied levels of electric stimulation.

## 2 Methods

### 2.1 Animals

This study utilized adult 8-12 week old heterozygous CX3CR1^+/GFP^ mice on a C57BL/6J background. An approximately equal mix of male and female animals were used. A subset of subjects were heterozygous for Thy1^+/ YFP^, in which spiral ganglion neurons express YFP. However, in an effort to maximize utilization of the available CX3CR1^+/GFP^ mice offspring, some subjects with a wildtype Thy1^+/+^ were used. No homozygous CX3CR1^GFP/GFP^ or Thy1^YFP/YFP^ subjects were used in this study. Genotyping was performed for CX3CR1 and Thy1 using standard PCR of genomic DNA from tail samples (Feng et al., 2000; Jung et al., 2000). Five experimental groups (n=3 per time-point) comprised this study: **Sham Surgery**, the approach to the round window niche was made, opening the round window but not inserting a CI; **Acute Insertion** (AI), the round window was opened and a CI briefly placed and removed; **Chronic Insertion** (ChI), full implantation of a CI without electric stimulation; **Low Stimulation** (LS), full CI implantation with low level electric stimulation starting on post-operative day 7; **High Stimulation** (HS), full CI implantation with high level electric stimulation starting on post-operative day 7. Surgery was performed exclusively on left ears through a round window approach with a custom 3 half-banded electrode CI (Cochlear Ltd., AUS), as previously described (Claussen et al., 2019). Mice were followed for varying timepoints until sacrifice and cochlear histology, including 4, 24 and 96 hours, 7, 11, 14, and 21 days post-operatively (**Figure 1** depicts the experimental timeline). The Low and High Stimulation groups were not included at the 4, 24 and 96 hour timepoints as this condition was identical to the chronic insertion group prior to electric stimulation start on day 7. The Sham Surgery and Acute Insertion groups were not followed at days 8,11 and 21 and day 21, respectively, as interim analysis showed no significant changes in cochlea histology at other timepoints. Three mice (n=3) were included at each timepoint for all groups. A separate control group (n=3) included mice who did not undergo surgery. The study protocol was approved by the University of Iowa Institutional Animal Care and Use Committee.

**Figure 1.**
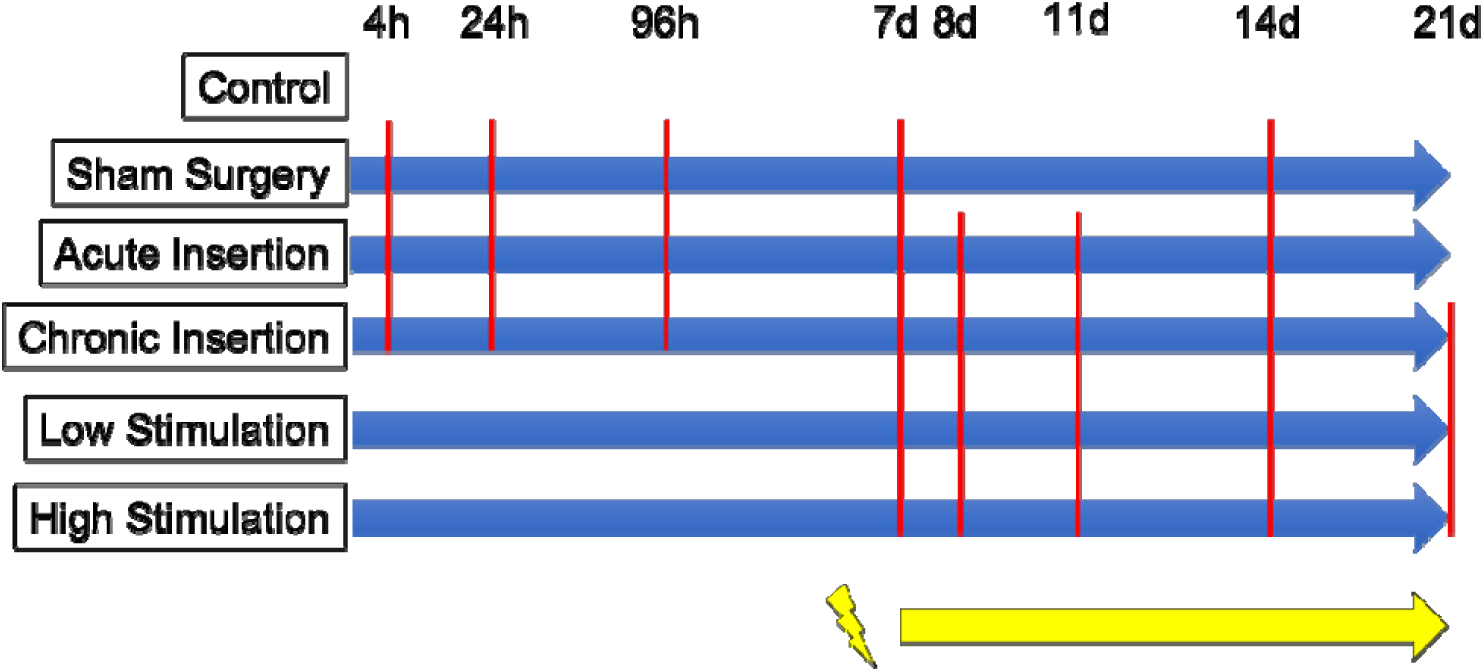
Experimental timeline. Timepoints are in reference to post-surgical time. Red lines mark separate timepoints included in respective groups. Electric stimulation in the relevant groups started on day 7, denoted by the yellow line.

### 2.2 Electric stimulation, impedance measurements and neural response telemetry (NRT)

Impedance measurements, NRT and electric stimulation programming were performed in Custom Sound EP 4.2 (Cochlear Ltd., AUS) in a procedure identical to that previously published (Claussen et al., 2019). Chronic Insertion, Low Stimulation and High Stimulation groups underwent Impedance and NRT threshold measurements for each separate electrode immediately prior to implantation, immediately post-operatively and at least weekly thereafter. Electric stimulation was performed by designing a MAP with a dynamic range of 1 between threshold and comfort levels. All functioning electrodes (impedance £ 35kOhms) were shorted together during stimulation using a software patch in Custom Sound EP 4.2 (Cochlear Ltd., AUS). This strategy allowed uniform electric stimulation while mice were connected to the CI processor in previously described stimulation cages for 5 hours per day, 5 days a week starting on post-operative day 7 (Claussen et al., 2019). The Low Stimulation and High Stimulation groups were programmed to a “threshold” and “comfort” level at 30CL below NRT threshold and behavioral response threshold, respectively. The MAP was re-adjusted weekly according to any changes in electrode functioning or NRT threshold measurement changes. Within the parameters of the custom CI and emulator system used, 0 CL corresponds to 17.5 μA and 0.44 nC/phase and 255 CL to 1750 μA and 43.75 nC/phase.

### 2.3 Histology

Left cochleae were harvested at the final respective timepoint for each subject and perfusion fixed with 4% paraformaldehyde. Cochleae were decalcified in 0.1M EDTA (pH 7.5) solution on a rotator that is changed regularly. Confirmation of the end point of decalcification was performed by combining equal parts of 5% Ammonium Hydroxide, 5% Ammonium Oxalate, and used decalcification solution from the specimen container. The decalcification process continued if the solution remained cloudy after 15 minutes. After decalcification, cochleae were washed three times for 10 minutes with PBS. Cochleae were then submersed in a cryoprotectant solution starting at 10% and increasing the cryoprotectant solution concentration by 10% every hour stopping at 30%. Cryoprotected cochleae were stored at -20 C until ready for sectioning. Cochleae were then infused with O.C.T. embedding medium (Tissue-TEK) and mounted to microtome stage with O.T.C. and dry ice. Using the sliding block microtome (American Optical 860) each cochlea was then sectioned in the mid-modiolar plane at 30 microns. Sections were then placed on slides for immunostaining and imaging. CX3CR1^+/GFP^ mice demonstrated endogenously green fluorescent monocytes and macrophages and Thy1^+/YFP^ mice exhibited endogenously yellow fluorescent spiral ganglion neurons and processes. When available, the YFP channel was included in histology pictures. Nuclear staining was performed with 1:1000 Hoechst (Hoechst 33342, Thermo Scientific).

### 2.4 Microscopy and macrophage quantification

A Leica SP5 (Leica, Germany) inverted confocal microscope was used to acquire 20x images of three serial sections per cochlea that included full views of the implanted basal turn. Imaris software (Bitplane, Switzerland) was used to perform volumetric cell counting through an entire histologic section z-stack. Regions of interest (ROI) including the Scala Tympani (ST), Lateral Wall (LW), Rosenthal’s Canal (RC) were defined by personnel familiar with cochlear microanatomy and blinded to experimental condition. Specifically, the LW region included both the spiral ligament and stria vascularis regions. The volume of the ROI as well as the overall number of all cell nuclei and GFP-positive cells were automatically quantified within the Imaris software and then manually verified by the examiner. These data were used to calculate the ratio of GFP-positive (macrophage) to total cell nuclei and density of GFP-positive cells (macrophages) in a defined ROI volume.

### 2.5 S*tatistical analysis*

Data were analyzed in GraphPad Prism 9 (Graphpad, USA). Graphs and histology images were made in Adobe Illustrator (Adobe, USA). Analysis of macrophage density and ratio were performed via a mixed effects model with group and time as independent variables, with follow-up multiple comparisons Tukey’s test when significant effects were encountered. Statistical significance was defined as p<0.05.

## 3 Results

### 3.1 Electrode impedance increases with time

**Figure 2** depicts impedance measurements overtime for separate electrodes at their pre-implantation, post-implantation and final timepoint measurements. Pre-implantation, conditioned electrode impedances ranged from 7.13-17.68 kOhms and post-implantation impedances ranged 9.13-23.85 kOhms (with one outlier at 37.22C kOhms that remained elevated above 35 kOhms on subsequent measurements). This study comprised 162 separate intracochlear electrodes in 54 cochlear implants. Most electrodes (147 or 90.7%) showed a gradual increase in impedance overtime, with a smaller portion of electrodes (15 or 9.3%) showing a sharp increase in impedance to the system testing limit of 125 kOhms, which denotes an open circuit and possible hardware failure (e.g. electrode lead wire fracture). These two distinct patterns of impedance creep are consistent with prior observations in this mouse CI model (Claussen et al., 2019). Within the experimental system, electrodes with impedances at or below 35 kOhms are considered functional; electrodes with impedances greater than 35 kOhms are unable to be stimulated as they are considered to exceed safe limits defined by the Shannon Equation. **Table 1** tabulates the number of functional electrodes on each implant at each respective endpoint (n=3 per timepoint and condition). At the furthest timepoint, 21 days post-implantation, 6/9 (66.7%) implants maintained at least 2 functional electrodes, enabling continued stimulation at this point. This is improved from prior, reporting only 25% of implants maintaining 2 functional electrode at 21 days post-implantation (Claussen et al., 2019). NRT thresholds were recorded weekly and ranged from 90 – 150CL across the cohort of all functional electrodes.

**Figure 2.**
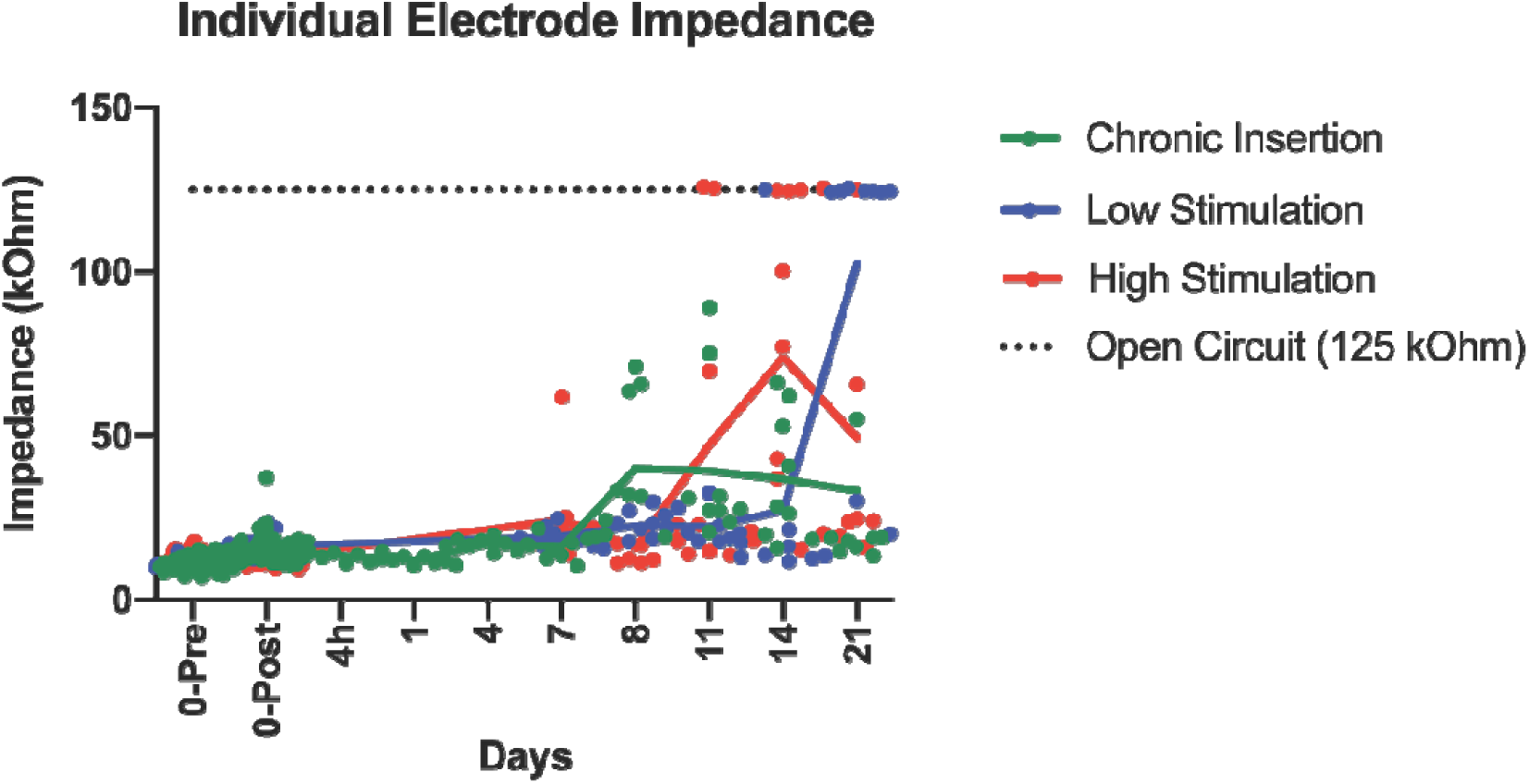
Baseline and final individual electrode impedance. The individual electrode impedance for the 3 intracochlear electrodes (E1,E2,E3) is plotted, including the initial values immediately before (0-Pre) and after (0-Post) implantation and final timepoint. The 125 kOhm value for an open circuit is plot as a dotted black line for reference.

**Table 1.** Final Electrode Functional Status. Number of functional electrodes per CI at the final time-point for the respective group and timepoint combinations. The number of functional electrodes per electrode at the final timepoint for each CI is listed, separating individual subjects with a “,”. A functional electrode was defined as having an impedance £ 35 kOhms.

| Number of functional ($\leq 35$ kOhms) electrodes per implant (n=3 per timepoint) | | | | | | | | |
| --- | --- | --- | --- | --- | --- | --- | --- | --- |
|  | 4 hours | 1 day | 4 days | 7 days | 8 days | 11 days | 14 days | 21 days |
| Chronic Insertion | 3,3,3 | 3,3,3 | 3,3,3 | 3,3,3 | 0,3,3 | 1,3,3 | 1,1,3 | 2,2,3 |
| Low Stimulation |  |  |  | 3,3,3 | 3,3,3 | 3,3,3 | 2,3,3 | 0,0,2 |
| High Stimulation |  |  |  | 2,3,3 | 3,3,3 | 1,3,3 | 0,1,1 | 0,3,3 |

### 3.2 Cochlear histologic changes following cochlear implantation

**Figure 3** depicts representative mid-modiolar sections across groups and timepoints. At baseline in control subjects, a small population of cochlear CX3CR1+ cells was seen throughout the cochlea, notably present within the modiolus, RC, LW and bone marrow spaces within the otic capsule, but absent from the ST. The majority of CX3CR1+ cells displayed a ramified morphology with dendritic processes. A varied increase in CX3CR1+ cells was noted across all experimental groups throughout the cochlea, prominently appearing in the LW, RC and modiolus and continuing to exhibit a majority ramified morphology. This influx of CX3CR1+ cells was most robust in the chronically implanted groups (ChI, LS and HS), peaking in the LW between days 7-21 and in RC between days 8-14.

**Figure 3.**
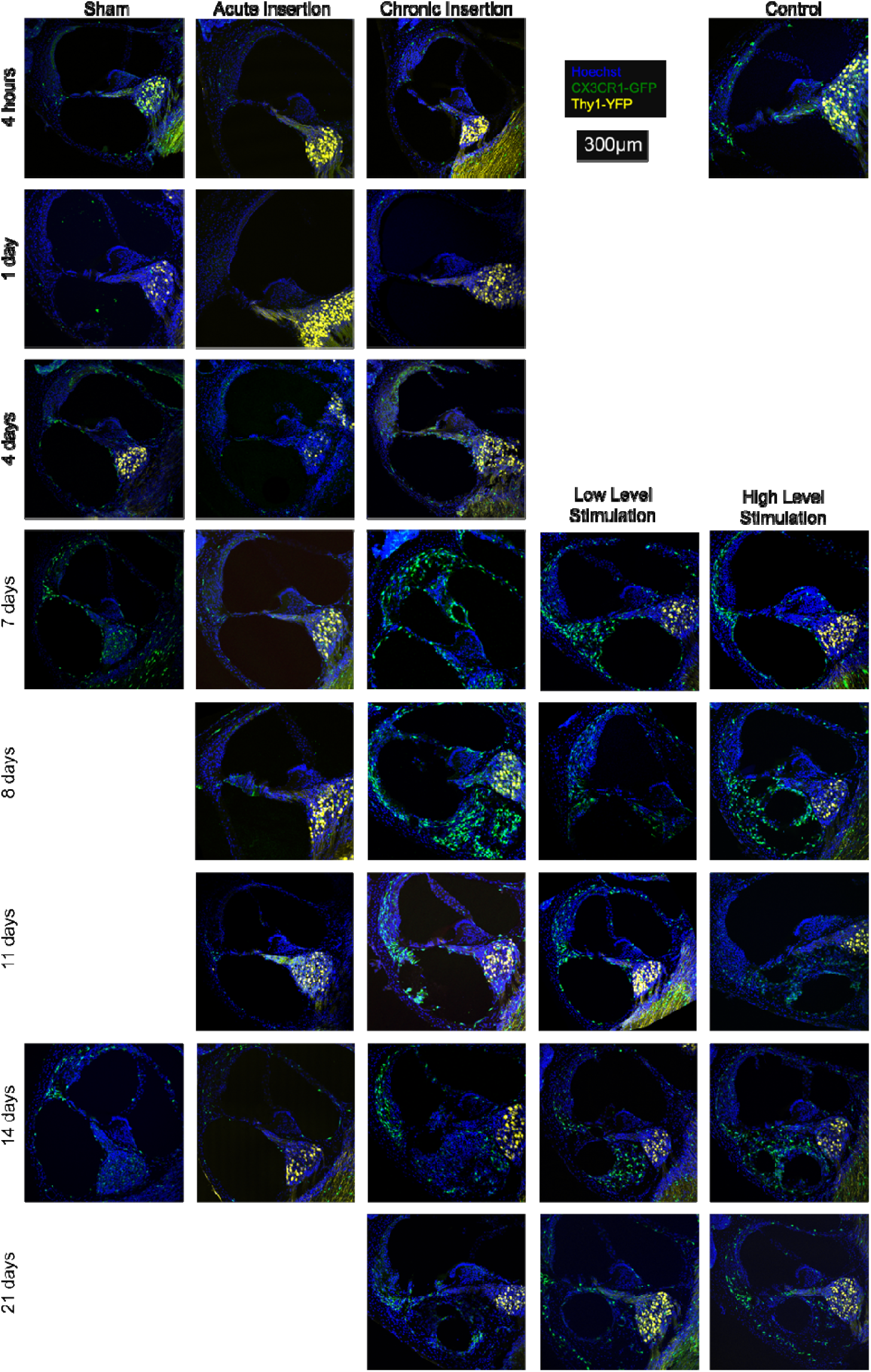
Cochlear fluorescence microscopy after chronic implantation. Mid-modiolar sections of the basal turn of the left cochlea from respective groups. Macrophage infiltration of the cochlea appears to increase over time and in the presence of the cochlear implant. Cellular infiltration into the ST was seen 7 days after implantation, accompanied by macrophage infiltration. Images show nuclei (Hoechst, blue), macrophages (CX3CR1-GFP, green) and neurons, (Thy1-YFP, yellow).

In response to chronic implantation (ChI, LS and HS groups), a robust cellular infiltrate, including CX3CR1+ cells, was visible in the ST starting on post-operative day 4. As seen in **Figure 3**, this tissue response formed around the CI electrode array and sometimes completely filled the ST space. The morphology of CX3CR1+ cells within the ST tissue response was more heterogeneous compared to the rest of the cochlea, often including diffusely fluorescent amoeboid cells alongside the ramified cells (**Figure 4)**. The tissue response extended from the round window to the basal turn of the ST but was not seen at areas estimated to be distal to the depth of CI insertion (middle and apical turns). Notably, the AI group displayed a cellular infiltrate immediately adjacent to the round window, mostly absent of CX3CR1+ cells, but this did not extend further into the basal turn of the ST in any subjects (**Figure 5**). No similar tissue response was seen at the round window in the sham or control groups. No obvious scalar translocations were seen amongst the AI, ChI, LS or HS groups.

**Figure 4.**
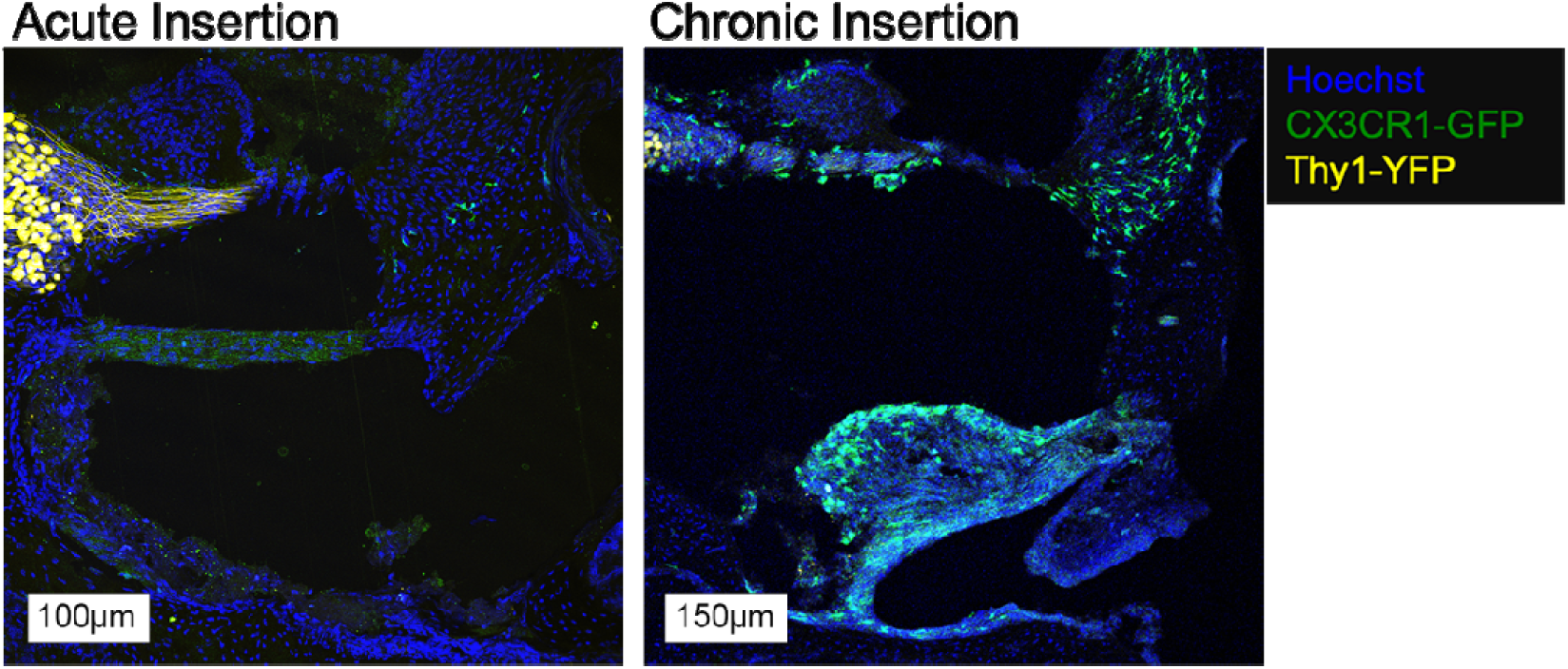
Round window membrane in AI and ChI mice at day 7. A small tissue response confined to the round window, without an implant tract is seen in the AI group with a small number of CX3CR1+ cells. A similar tissue response near the round window with an implant tract is seen in the ChI group, accompanied by a more robust monocyte/macrophage infiltrate. Images show cell nuclei (Hoechst, blue), macrophages (CX3CR1-GFP, green) and neurons, (Thy1-YFP, yellow).

**Figure 5.**
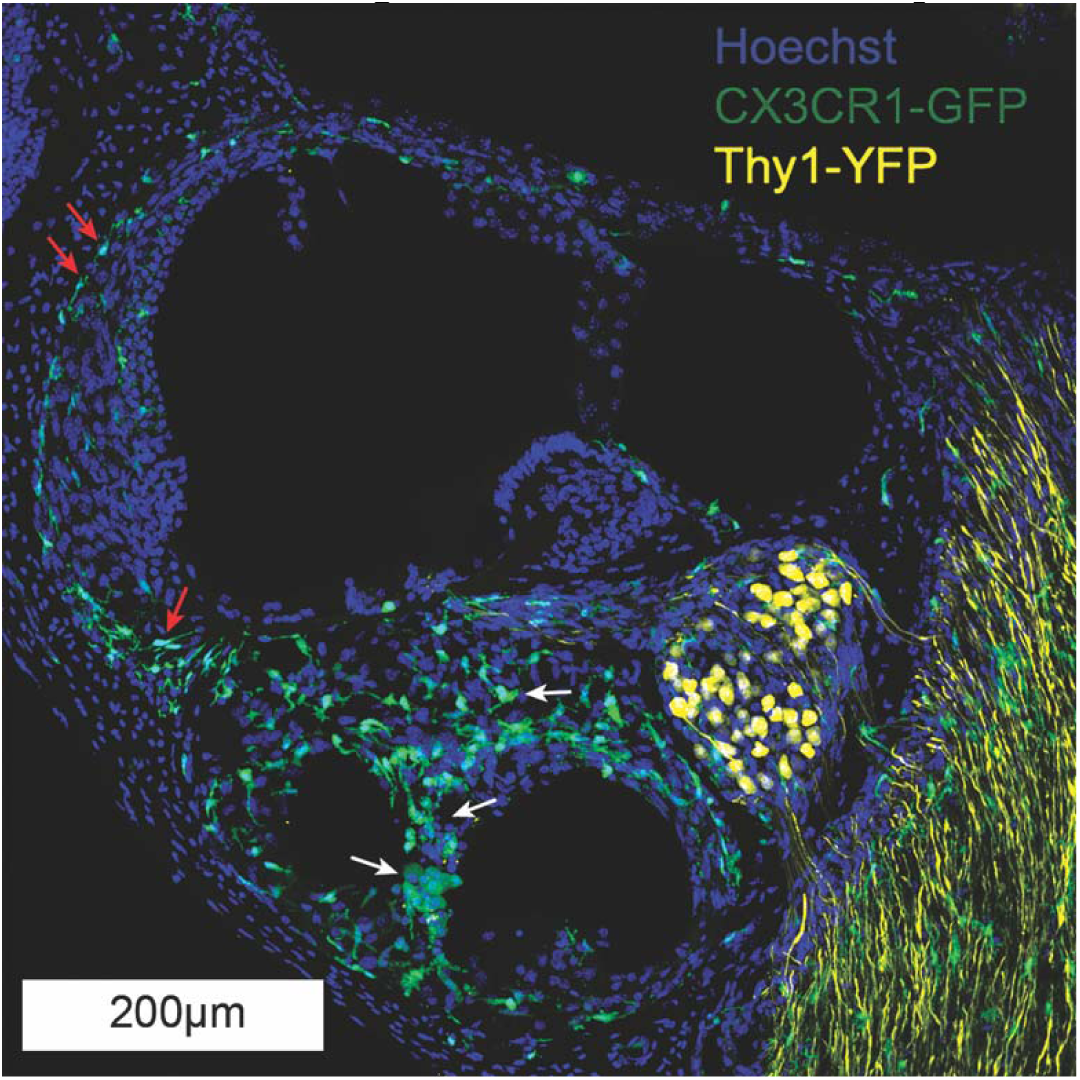
Basal turn, 14 day High Stimulation. Red arrows highlight the ramified macrophages and white arrows the ameboid macrophages, located in fibrotic tissue of the scala tympani. Labeling includes nuclei (Hoechst, blue), macrophages (CX3CR1-GFP, green) and neurons, (Thy1-YFP, yellow).

### 3.3 Quantification of CX3CR1+ cellular infiltration

#### 3.3.1 Rosenthal’s canal

**Figure 6** depicts quantification of CX3CR1+ cell density and ratio of CX_3_CR1+ cells to total cell nuclei in Rosenthal’s Canal. CX3CR1+ cell infiltration was minimal among all groups until post-operative day 4, when both CX3CR1+ cell density and ratio raised in both the ChI and Sham groups. CX3CR1+ cell infiltration was generally seen to be elevated in the ChI, LS and HS groups compared to all other groups, reaching a peak from post-operative day 8 to 14 and falling slightly at post-operative day 21. The sham and AI groups showed peaks in CX3CR1+ cell quantification at post-operative day 4 and 7, respectively, but maintained values close to controls at other time-points. CX3CR1+ cell density and ratio were significantly (p<0.05) increased in the ChI and LS groups compared to controls from post-operative day 7-21. CX3CR1+ cell ratio was significantly increased (p<0.05) in the ChI, LS and HS groups compared to controls and acute insertion from post-operative day 7-21. There was no significant difference amongst chronically implanted groups at any time-point. As previously mentioned, YFP neuronal labelling was not available in all subjects, thus we are unable to make comparisons among conditions regarding spiral ganglion neuron survival and response after cochlear implantation.

**Figure 6.**
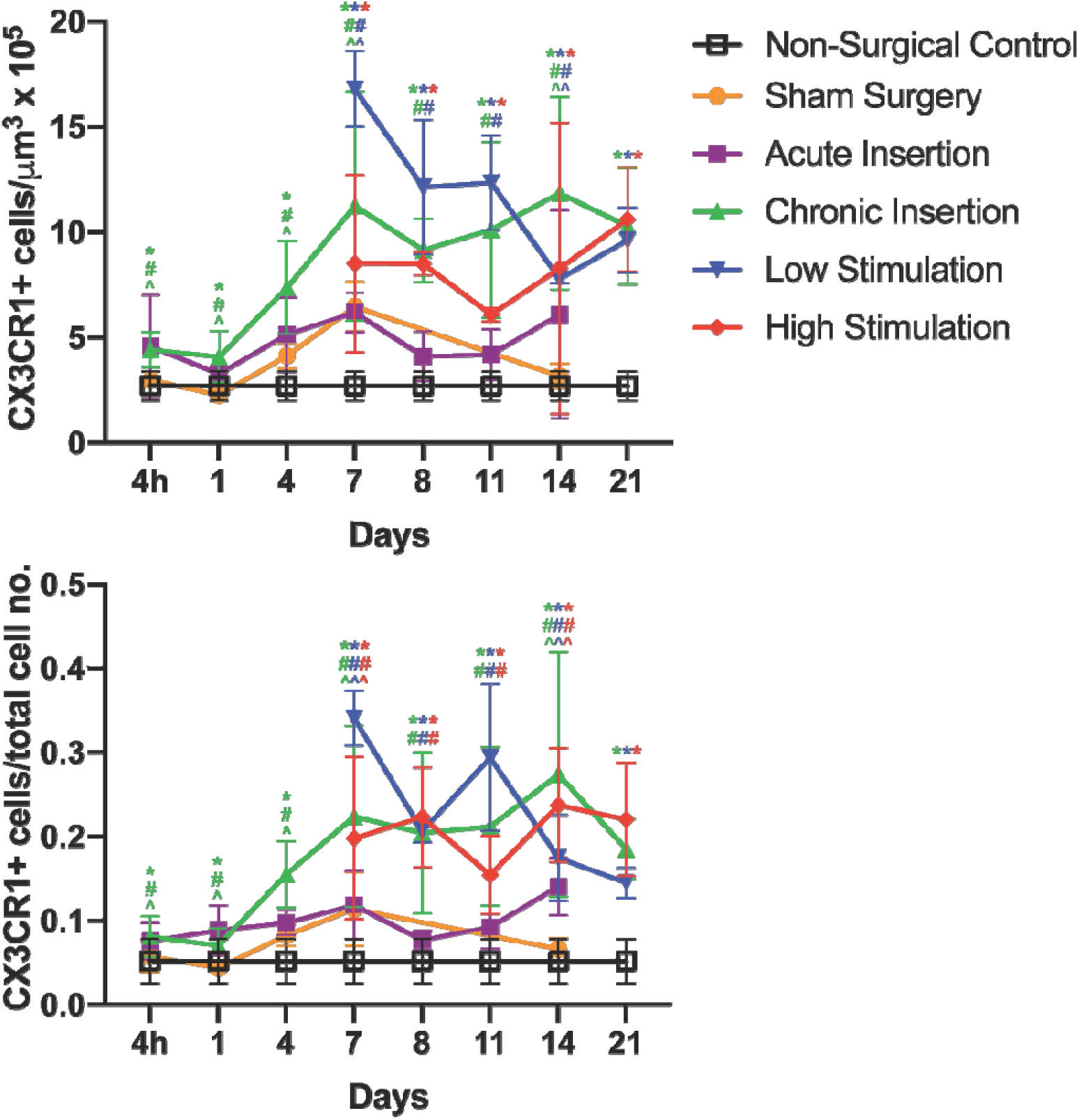
CX3CR1+ cell density and ratio in the lateral wall. Density was calculated as quantity of CX3CR1+ cells per unit volume. Ratio was calculated as quantity of CX3CR1+ cell per total number of cells. “*”, “#” and “^” are color coded to respective groups to denote statistically significant differences (p<0.05) compared to the control, AI and sham surgery groups, respectively. Error bars represent standard deviation.

#### 3.3.2 Lateral wall

Quantification of CX3CR1+ cell density and ratio to total cell nuclei in the lateral wall region (including the stria vascularis and spiral ligament) is depicted in **Figure 7**. Elevated CX3CR1+ cell infiltration was seen in both the ChI and AI groups as early as 4 hours post-implantation. The AI group maintained a steady plateau of increased CX3CR1+ cell density and cell number compared to controls across all time-points, whereas the ChI group showed continued increase in CX3CR1+ cell infiltration until reaching a steady state from day 7 to 21. Similarly, the CX3CR1+ cell density and ratio remained elevated in the LS and HS groups compared to controls from post-operative day 7 to 21, with a notable peak in LS on day 7. The ChI, LS and HS groups showed significantly (p<0.05) greater CX3CR1+ cell density and ratio compared to controls, AI and sham across timepoints. There were no significant (p<0.05) differences between ChI, LS and HS groups at any timepoint.

**Figure 7.**
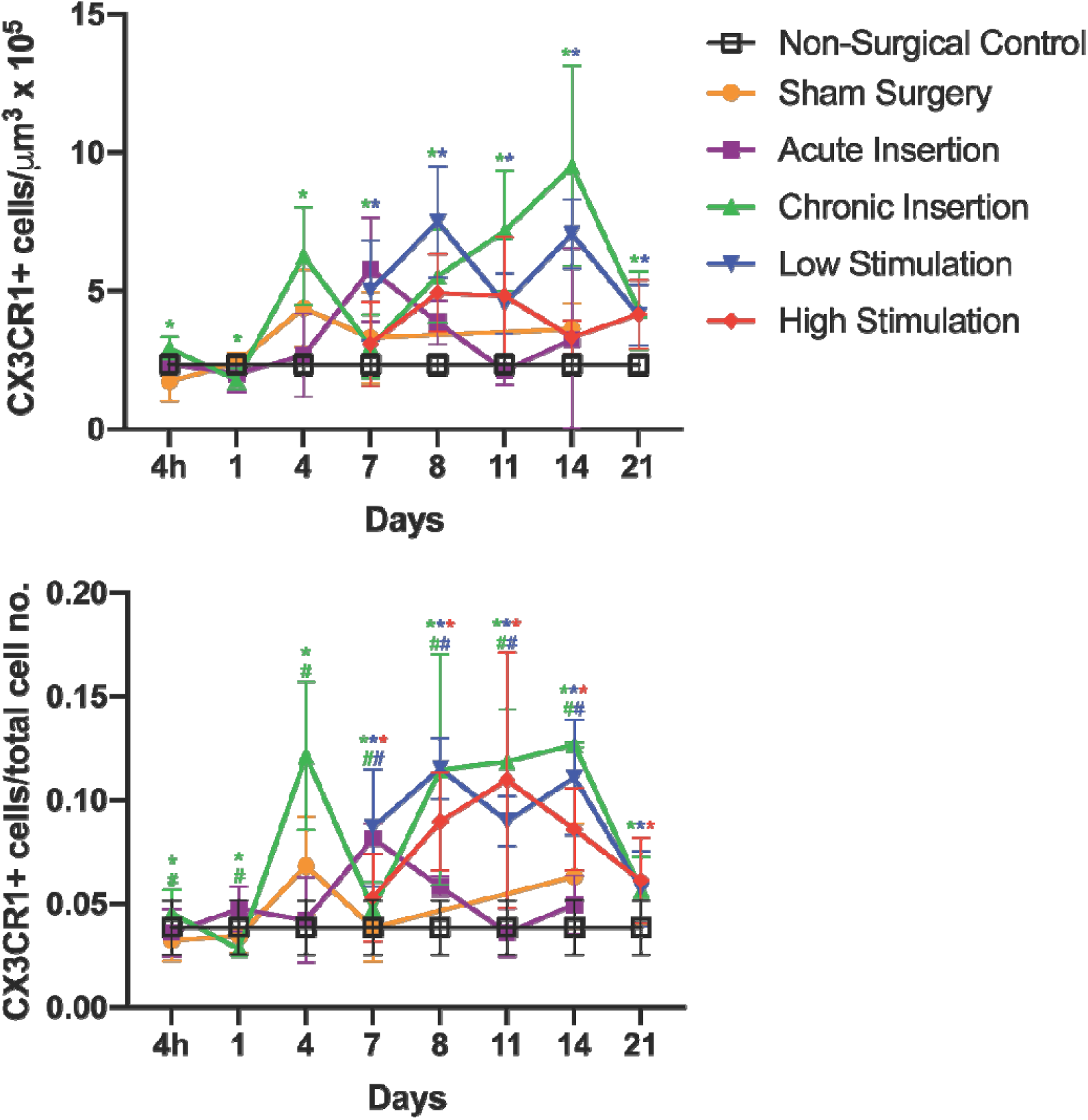
CX3CR1+ cell density and ratio in Rosenthal’s Canal. Density was calculated as quantity of CX3CR1+ cells per unit volume. Ratio was calculated as quantity of CX3CR1+ positive cell per total number of cells. “*” and “#” are color coded to respective groups to denote statistically significant differences (p<0.05) compared to the control and AI and groups, respectively. Error bars represent standard deviation.

#### 3.3.3 Scala tympani

**Figure 8** depicts the quantification of CX3CR1+cell density and ratio in the ST. A ST tissue response, including CX3CR1+ cell infiltration and increased cellularity, was seen as early as 4 days post-implantation in the ChI group, with all implanted subjects (ChI, LS and HS groups) showing an intrascalar cellular infiltration comprised of both CX3CR1+cells and other GFP-negative cells by post-implantation day 7, which persisted until the final time-point at day 21. CX3CR1+ cell density and ratio in the 3 implanted groups was significantly (p<0.05) greater compared to control, AI and Sham from day 4 onward. There was no significant (p<0.05) difference between the ChI, LS and HS groups. CX3CR1+ cell density remained persistently elevated in the implanted groups until the final timepoint at day 21, however CX3CR1+ cell ratio showed a consistent decline amongst all 3 chronically implanted groups on day 21. This reduction in CX3CR1+ cell ratio in the setting of a steady CX3CR1+ cell density suggests an accumulation of other GFP-negative immune (e.g. leukocytes) or non-immune cells (e.g. fibroblasts) within the ST at this final time-point.

**Figure 8.**
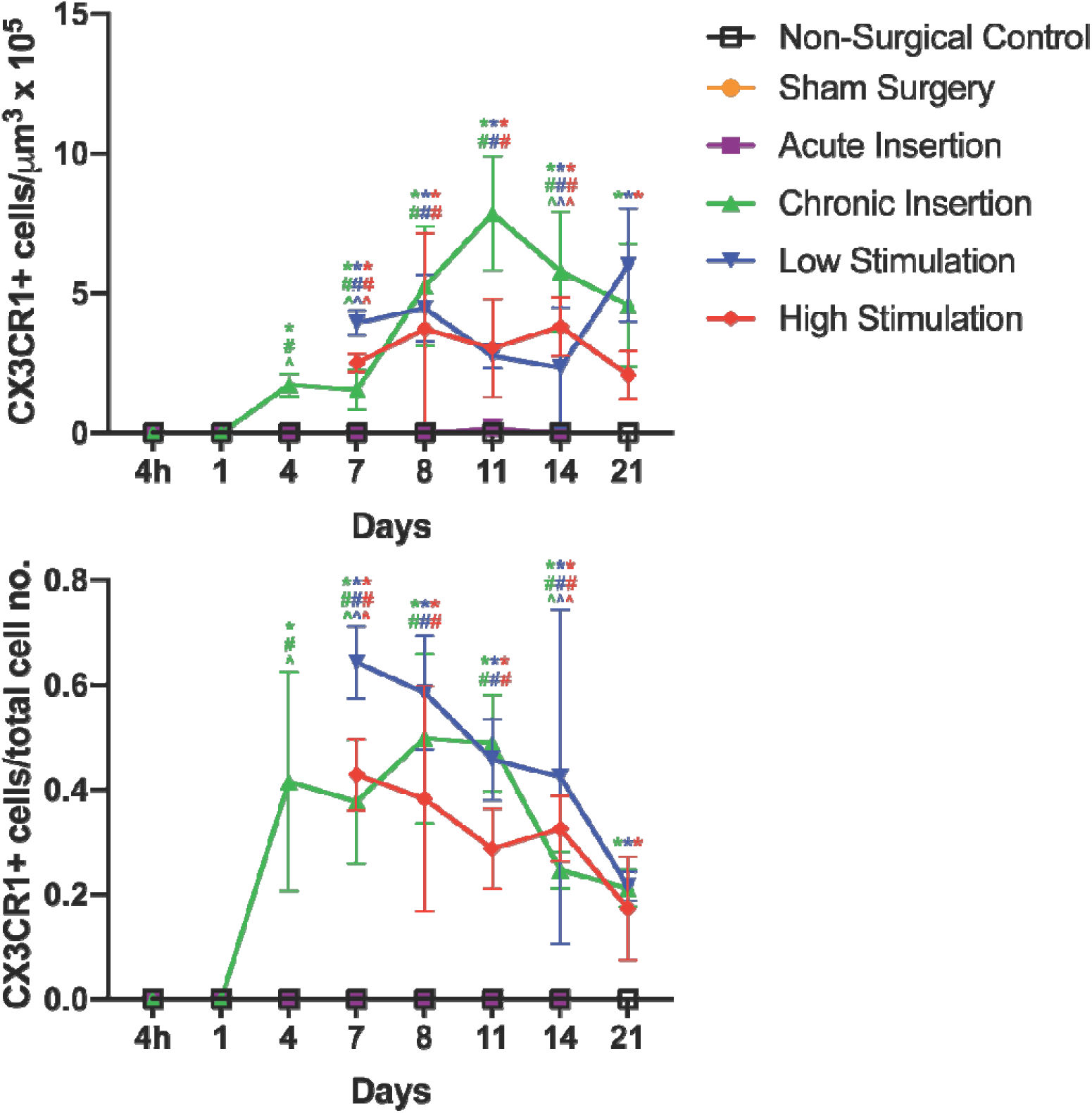
CX3CR1+ cell density and ratio in the Scala Tympani. Density was calculated as quantity of CX3CR1+ cells per unit volume. Ratio was calculated as quantity of CX3CR1+ cells per total number of cells. “*”, “#” and “^” are color coded to respective groups to denote statistically significant differences (p<0.05) compared to the control, AI and sham surgery groups, respectively. Error bars represent standard deviation.

## 4 Discussion

This study was designed to examine the cochlear innate immune response to cochlear implantation in CX3CR1^+/GFP^ mice across multiple timepoints and assess the separate contributions of surgical trauma (control vs sham vs AI vs ChI), presence of the CI foreign body (AI vs ChI) and electric stimulation (ChI vs LS vs HS). Past work suggests cochlear CX3CR1+cells primarily represent a population of resident cochlear macrophages in both normal and injured cochleae, enabling assessment of the cochlear innate immune response in this study (Hirose et al., 2005). Although the number of animals per group was limited secondary to the high number of group and timepoint combinations, several consistent patterns of response were observed. As expected from prior studies, a baseline cohort of CX3CR1+ cells was seen throughout the cochlea, notably absent from the acellular spaces of the scalae. Any surgical trauma, including round window opening (sham) and acute CI insertion (AI) caused an increase in CX3CR1+ cells within the lateral wall and Rosenthal’s canal compared to baseline control, which was more pronounced with chronic CI insertion (ChI, LS and HS). Surgical trauma alone was not sufficient to induce a ST tissue response beyond the round window; Chronic CI insertion was necessary for formation of a ST tissue response. Varying the degree of electric stimulation did not affect the innate immune response to cochlear implantation. Taken together, these observations suggest surgical trauma alone may induce a cochlear innate immune response, but the chronic presence of an indwelling electrode array further augments the cochlear inflammatory response and is necessary in formation of a ST tissue response. Although likely multifactorial, this observation points to a critical role that the foreign body response to cochlear implant biomaterials plays in the cochlear inflammation, fibrosis and neo-ossification following CI.

The robust peri-implant cellular infiltrate in the ST of the chronically implanted groups was similar in extent to that previously published in the mouse CI model, in which the ST was occupied with fibrotic tissue and areas of neo-ossification confined to the depth of CI insertion. (Claussen et al., 2019). CX3CR1+ cells were seen to comprise a substantial element of the ST response between days 7 and 11, accounting for over 50% of the cells within the ST of some subjects. Similar patterns of tissue response have been documented in implanted human temporal bones, accompanied by both innate (e.g. monocytes, macrophages) (Noonan et al., 2020; Okayasu et al., 2020) and adaptive immune cells (e.g. B-cells, T-cells) (Nadol et al., 2014). However, the human intrascalar tissue response is sometimes seen to extend further past the depth of CI insertion or into other scala, especially in the setting of scalar translocations (Kamakura & Nadol, 2016; Li et al., 2007). The current key findings of the tissue response being exclusively present in the chronically implanted groups and confined to areas directly adjacent to the CI highlights the role of the electrode array itself in driving and organizing the tissue response. Evidence of such a foreign body response has been seen in humans in the form of phagocytosed implant materials and foreign body giant cells (Nadol et al., 2008; O’Malley et al., 2017) and was recently reviewed by Foggia et al. (Foggia et al., 2019). In this special issue, Jensen et al., present new data comparing the foreign body response to different CI materials (e.g. silastic and platinum).

It is not understood what effect, if any, electric stimulation may have on the cochlear immune response and fibrosis after cochlear implantation and if it would be mediated by electrode material composition or direct electric effect on surrounding tissues. Different patterns of tissue fibrosis surrounding the CI relating to the location of the platinum electrodes have been seen in human temporal bones (Ishai et al., 2017). Further, platinum dissolution into the scala has been demonstrated in guinea pig models of CI at charge densities above those used in the clinical setting, with the potential consequence of enhancing the foreign body response (Shepherd et al., 2019). We did not find any effect of the levels of electric stimulation used in this study on the innate immune response or degree of ST cellular infiltrate. Although overall rates of hardware failure (e.g. electrode impedance increasing to a level prohibiting electric stimulation) improved in this study prior to that published in Claussen et al. (2019), impedance creep and electrode failure did occur, hampering electric stimulation at the longest time points in the LS group, in particular. Notably, the charge densities used in this study were approximately 2 orders of magnitude below those shown to result in local platinum dissolution from the CI electrodes. Further, the robust and rapid development of a ST cellular infiltrate in the mouse CI model may obscure any small effects resulting from varied electric stimulation intensities, such as distribution of soft tissue fibrosis versus neo-ossification, which were not detectable with the histologic methods used in this study. We refer readers to Jensen et al. in this issue, where 3D X-ray microscopy was used to demonstrate a greater propensity for neo-ossification around platinum surfaces of the CI as opposed to silicone surfaces in a mouse CI model.

In the current study, we cannot completely separate the role of insertion trauma from the foreign body effects of an indwelling CI toward eliciting a cochlear innate immune response, as the potential for insertion trauma is inherent to chronic cochlear implantation. However, round window opening and acute CI insertion alone, in the absence of obvious scalar translocations, was not sufficient to generate a tissue response beyond the immediate round window, but did elicit a cochlear innate immune response, albeit to a lesser degree, to chronic implantation. Other factors shown to promote cochlear fibrosis include surgical trauma from cochleostomy, use of muscle to seal round window and perilymphatic introduction of bone dust (McELVEEN et al., 1995; Rowe et al., 2016). However, these factors may not have influenced this study, as a round window approach was used that involved minimal drilling of the round window niche with irrigation of any bone dust prior to round window opening and use of fascia, not muscle to seal the round window. A cellular infiltrate mostly devoid of CX3CR1+ cells was seen immediately at the round window in the acute insertion group, which may relate to the local irritation from round window niche drilling or the placement of a fascia graft to seal the round window. There is ample evidence to suggest that major insertional trauma events, including scalar translocation and osseus spiral lamina fracture, may influence fibrosis and neo-ossification after cochlear implantation, however none of these events were seen in the present study to validate these causative factors (Kamakura & Nadol, 2016; Li et al., 2007; O’Leary et al., 2013). Based on these present and past findings, we observe that the CI insertion event generates an innate immune response and speculate that the immune cells and accompanying milieu of inflammatory mediators (e.g. chemokines, cytokines, growth factors) help initiate the foreign body response and fibrotic tissue reaction to the CI in an exposure dependent manner that is commensurate with the degree of insertional trauma. Understanding the mechanisms of how insertion trauma elicits an innate immune response is beyond the scope this study, but we hypothesize this broadly includes initiation of sterile inflammation through direct (e.g. physical cellular insult from force of insertion) and collateral (e.g. altered homeostasis and cell death related to trauma) generation of damage-associated molecular patterns and other inflammatory signals, the degree to which may vary with the severity of insertion trauma. (Klegeris, 2021; Mariani et al., 2019; Wood & Zuo, 2017).

In the normal mouse cochlea, CX3CR1+ cells mostly represent resident cochlear macrophages along with a smaller subset of other immune cells (e.g. NK cells, T cells) (Hirose et al., 2005). The increase in CX3CR1+ cells in the current study, especially in the chronic insertion groups, is similar to that following other cochlear injuries, including noise exposure and isolated outer hair cell ablation, and is seen to consistently peak 14 days post-injury across several studies (Kaur et al., 2015; Rai et al., 2020; Tan et al., 2016). The nature of this increase in cochlear CX3CR1+ cells is likely due to migration of cochlear macrophages from the circulation, based on prior studies examining local proliferation of CX3CR1+ cells in the cochlea after noise (Hirose et al., 2005). An increase in circulating monocytes and tissue macrophages that exhibit low or no expression of CX3CR1+ also occurs following cochlear, neural and other tissue injury, however, the current study was not designed to detect these additional elements of the innate immune response (Hirose et al., 2014; Puntambekar et al., 2018; Wynn & Vannella, 2016). As evidence of a continued evolution of the inflammatory response, we saw a decrease in the ratio of macrophages to total number of cells within the ST in the chronically implanted groups from day 8 to 21 in the setting of a relatively stable trend of CX3CR1+ cell density. This finding may provide indirect evidence of evolving macrophage phenotype away from CX3CR1 + cells and an increase in other cells involved in fibrosis (e.g. myofibroblasts) that would contribute to maturation of the inflammatory response. The nature of cochlear implantation involving a chronic insult in the form of the CI foreign body may produce diverging temporal courses and phenotypes of the immune response compared to the other studies mentioned, which involve acute, single event injuries. The current study is limited by the absence of hearing loss prior to cochlear implantation and any associated alterations to or priming of the baseline resident macrophage population. Future work is needed to more broadly examine the evolving macrophage phenotypes not captured in the CX3CR1^+/GFP^ mouse CI model and to include prior deafening to more closely model the cochlear immune state prior to implantation as seen in the clinical context.

CX3CR1+ macrophages play diverse roles including local surveillance for pathogens, debris clearance of damaged cells, and antigen presentation. Macrophage morphology is often associated with function, with ramified cells playing a surveillance role and amoeboid cells representing an activated, phagocytic phenotype (Savage et al., 2019). In the present study, a diversity of amoeboid and ramified morphologies of CX3CR1+ cells were seen, which is consistent with findings in implanted human cadaveric studies (Noonan et al., 2020).

Macrophage phenotype or “polarization” has further been characterized along a spectrum of M1 (inflammatory) and M2 (anti-inflammatory) subtypes. Recent transcriptomic and flow cytometry data report additional complexity in distinguishing subtypes and their contributions to inflammation, fibrosis, tissue repair and regeneration (Parakalan et al., 2012; Rai et al., 2020; Wynn & Vannella, 2016). Within the cochlea, CX3CR1+ macrophages have been implicated in protective and homeostatic functions for spiral ganglion neurons following hair cell or direct neuronal injury (Kaur et al., 2015; Lang et al., 2016). Conversely, in other tissues, CX3CR1+ macrophages have shown an inflammatory and pro-fibrotic phenotype, particularly in the setting of pulmonary fibrosis (Aran et al., 2019; Wynn & Vannella, 2016). This pro-fibrotic subtype may be relevant to the present CI model, as the most robust increase in CX3CR1+ cells was seen in the fibrous tissue growth around the CI. However, the current data cannot definitively attribute any pro-fibrotic or other specific role CX3CR1+ macrophages in the cochlea.

These experiments highlight the utility of a mouse model of functional CI, which can be combined with pre-existing transgenic mouse models to investigate both the cochlear immune response to CI (as in this study) and potentially extend to investigating CI in other models of hearing loss. The applications CX3CR1^+/GFP^ mice in a CI model extends beyond direct observation of the cochlear innate immune response and may be useful for studying the efficacy and mechanism of future strategies directed at mitigating the post-implantation fibrotic response and preserving residual acoustic hearing in hybrid cochlear implantation. Limitations of the mouse CI model include CI lead wire fracture and hardware failure that limit long-term stimulation. However, the current experiments show improvement in rates of electrode preservation compared to prior work. The rapid time-course (7-8 days) of peri-implant ST tissue response in implanted mice may not accurately model the similar time-course seen in humans, but does result in a similar macrophage infiltrate and areas of fibrotic soft tissue and neo-ossification (Noonan et al., 2020; Seyyedi & Nadol, 2014).

## 5 Conclusion

Chronic CI placement in CX3CR1^+/GFP^ mice results in a robust innate immune response shown by an increase in CX3CR1 cells throughout the cochlea, accompanied by a peri-implant ST cellular infiltrate similar to that seen in humans. This tissue response is not significantly affected by the level of electric stimulation. Surgical trauma related to cochlear implantation alone initiated an innate immune response in the lateral wall and Rosenthal’s canal that was blunted compared to the response in chronically implanted cochleae. Further, surgical trauma alone did not lead to significant cellular infiltration into the ST. This suggests that the foreign body reaction to a chronically implanted device is more important than the contribution of surgical trauma in the formation of cochlear fibrosis and remodeling of the cochlear architecture. Future work is needed to understand the role CX3CR1+ macrophages in the cochlear response to cochlear implantation as well as further elucidate the broader innate and adaptive immune response over time. Such insights may be useful in developing and measuring the efficacy of strategies to reduce the cochlear tissue response and loss of residual acoustic hearing after cochlear implantation.

## Abbreviations

AI: acute insertion
ChI: chronic insertion
CI: cochlear implant
CL: clinical level
FBR: foreign body response
GFP: green fluorescent protein
HS: high stimulation
LS: low stimulation
LW: lateral wall
NRT: neural response telemetry
PBS: phosphate buffered saline
ROI: region of interest
RC: Rosenthal’s canal
ST: scala tympani
YFP: yellow fluorescent protein

## 6 Acknowledgements

Support for these studies was provided by National Institutes of Health, National Institute on Deafness and Other Communication Disorders, Besthesda MD [grant numbers R01 DC012578 (MRH), R01 DC011315 (KH), T32 DC000040 (ADC, MRH), and UL1TR002537 (MRH)], and Cochlear Americas, Centennial, CO [grant number 17881800-01].

## References

Aran, D., Looney, A. P., Liu, L., Wu, E., Fong, V., Hsu, A., Chak, S., Naikawadi, R. P., Wolters, P. J., Abate, A. R., Butte, A. J., & Bhattacharya, M. (2019). Reference-based analysis of lung single-cell sequencing reveals a transitional profibrotic macrophage. Nature Immunology, 20(2), 163–172. https://doi.org/10.1038/s41590-018-0276-y

Clark, G. M., Kranz, H. G., Minas, H., & Nathar, J. M. (1975). Histopathological findings in cochlear implants in cats. The Journal of Laryngology & Otology, 89(5), 495–504. https://doi.org/10.1017/S002221510008066X

Claussen, A. D., Quevedo, R. V., Mostaert, B., Kirk, J. R., Dueck, W. F., & Hansen, M. R. (2019). A mouse model of cochlear implantation with chronic electric stimulation. PLoS ONE, 14(4). https://doi.org/10.1371/journal.pone.0215407

Feng, G., Mellor, R. H., Bernstein, M., Keller-Peck, C., Nguyen, Q. T., Wallace, M., Nerbonne, J. M., Lichtman, J. W., & Sanes, J. R. (2000). Imaging neuronal subsets in transgenic mice expressing multiple spectral variants of GFP. Neuron, 28(1), 41–51. https://doi.org/10.1016/S0896-6273(00)00084-2

Foggia, M. J., Quevedo, R. V., & Hansen, M. R. (2019). Intracochlear fibrosis and the foreign body response to cochlear implant biomaterials. Laryngoscope Investigative Otolaryngology. https://doi.org/10.1002/lio2.329

Goman, A. M., Dunn, C. C., Gantz, B. J., & Lin, F. R. (2018). PREVALENCE of POTENTIAL HYBRID and CONVENTIONAL COCHLEAR IMPLANT CANDIDATES BASED on AUDIOMETRIC PROFILE. In Otology and Neurotology. https://doi.org/10.1097/MAO.0000000000001728

Hirose, K., Discolo, C. M., Keasler, J. R., & Ransohoff, R. (2005). Mononuclear phagocytes migrate into the murine cochlea after acoustic trauma. Journal of Comparative Neurology, 489(2), 180–194. https://doi.org/10.1002/cne.20619

Hirose, K., Li, S. Z., Ohlemiller, K. K., & Ransohoff, R. M. (2014). Systemic lipopolysaccharide induces cochlear inflammation and exacerbates the synergistic ototoxicity of kanamycin and furosemide. JARO - Journal of the Association for Research in Otolaryngology, 15(4), 555–570. https://doi.org/10.1007/s10162-014-0458-8

Ishai, R., Herrmann, B. S., Nadol, J. B., & Quesnel, A. M. (2017). The pattern and degree of capsular fibrous sheaths surrounding cochlear electrode arrays. Hearing Research, 348, 44–53. https://doi.org/10.1016/j.heares.2017.02.012

Jung, S., Aliberti, J., Graemmel, P., Sunshine, M. J., Kreutzberg, G. W., Sher, A., & Littman, D. R. (2000). Analysis of Fractalkine Receptor CX3CR1 Function by Targeted Deletion and Green Fluorescent Protein Reporter Gene Insertion. Molecular and Cellular Biology, 20(11), 4106–4114. https://doi.org/10.1128/mcb.20.11.4106-4114.2000

Kamakura, T., & Nadol, J. B. (2016). Correlation between word recognition score and intracochlear new bone and fibrous tissue after cochlear implantation in the human. Hearing Research, 339, 132–141. https://doi.org/10.1016/j.heares.2016.06.015

Kaufmann, C. R., Tejani, V. D., Fredericks, D. C., Henslee, A. M., Sun, D. Q., Abbas, P. J., & Hansen, M. R. (2020). Pilot Evaluation of Sheep as in Vivo Model for Cochlear Implantation. Otology and Neurotology. https://doi.org/10.1097/MAO.0000000000002587

Kaur, T., Zamani, D., Tong, L., Rube, E. W., Ohlemiller, K. K., Hirose, K., & Warchol, M. E. (2015). Fractalkine signaling regulates macrophage recruitment into the cochlea and promotes the survival of spiral ganglion neurons after selective hair cell lesion. Journal of Neuroscience, 35(45), 15050–15061. https://doi.org/10.1523/JNEUROSCI.2325-15.2015

Klegeris, A. (2021). Regulation of neuroimmune processes by damage- and resolution-associated molecular patterns. 16(3), 423–429.

Lang, H., Nishimoto, E., Xing, Y., Brown, L. S. N., Noble, K. V., Barth, J. L., LaRue, A. C., Ando, K., & Schulte, B. A. (2016). Contributions of mouse and human hematopoietic cells to remodeling of the adult auditory nerve after neuron loss. Molecular Therapy, 24(11), 2000–2011. https://doi.org/10.1038/mt.2016.174

Li, P. M. M. C., Somdas, M. A., Eddington, D. K., & Nadol, J. B. (2007). Analysis of intracochlear new bone and fibrous tissue formation in human subjects with cochlear implants. Annals of Otology, Rhinology and Laryngology, 116(10), 731–738. https://doi.org/10.1177/000348940711601004

Linthicum, F. H., Doherty, J. K., Lopez, I. A., & Ishiyama, A. (2017). Cochlear implant histopathology. World Journal of Otorhinolaryngology - Head and Neck Surgery, 3(4), 211–213. https://doi.org/10.1016/j.wjorl.2017.12.008

Mariani, E., Lisignoli, G., Borzì, R. M., & Pulsatelli, L. (2019). Biomaterials: Foreign bodies or tuners for the immune response? International Journal of Molecular Sciences, 20(3). https://doi.org/10.3390/ijms20030636

Mcelveen, J. T., Wolford, R. D., & Miyamoto, R. T. (1995). Implications of bone pâté in cochlear implant surgery. Otolaryngology - Head and Neck Surgery. https://doi.org/10.1016/S0194-5998(95)70284-9

Mitchell-Innes, A., Saeed, S. R., & Irving, R. (2018). The future of cochlear implant design. Advances in Oto-Rhino-Laryngology, 81, 105–113. https://doi.org/10.1159/000485540

Nadol, J. B., Eddington, D. K., & Burgess, B. J. (2008). Foreign Body or Hypersensitivity Granuloma of the Inner Ear After Cochlear Implantation: One Possible Cause of a Soft Failure? Otology and Neurotology, 29(8), 1076–1084. https://doi.org/10.1097/MAO.0b013e31818c33cf

Nadol, J. B., O’Malley, J. T., Burgess, B. J., & Galler, D. (2014). Cellular immunologic responses to cochlear implantation in the human. Hearing Research, 318, 11–17. https://doi.org/10.1016/j.heares.2014.09.007

Needham, K., Stathopoulos, D., Newbold, C., Leavens, J., Risi, F., Manouchehri, S., Durmo, I., & Cowan, R. (2020). Electrode impedance changes after implantation of a dexamethasone-eluting intracochlear array. Cochlear Implants International, 21(2), 98–109. https://doi.org/10.1080/14670100.2019.1680167

Noonan, K. Y., Lopez, I. A., Ishiyama, G., & Ishiyama, A. (2020). Immune Response of Macrophage Population to Cochlear Implantation: Cochlea Immune Cells. Otology & Neurotology□: Official Publication of the American Otological Society, American Neurotology Society [and] European Academy of Otology and Neurotology. https://doi.org/10.1097/MAO.0000000000002764

O’Leary, S. J., Monksfield, P., Kel, G., Connolly, T., Souter, M. A., Chang, A., Marovic, P., O’Leary, J. S., Richardson, R., & Eastwood, H. (2013). Relations between cochlear histopathology and hearing loss in experimental cochlear implantation. Hearing Research, 298, 27–35. https://doi.org/10.1016/j.heares.2013.01.012

O’Malley, J. T., Burgess, B. J., Galler, D., & Nadol, J. B. (2017). Foreign body response to silicone in cochlear implant electrodes in the human. Otology and Neurotology, 38(7), 970–977. https://doi.org/10.1097/MAO.0000000000001454

Okayasu, T., O’Malley, J. T., & Nadol, J. B. (2019). Density of macrophages immunostained with anti-iba1 antibody in the vestibular endorgans after cochlear implantation in the human. Otology and Neurotology, 40(8), E774–E781. https://doi.org/10.1097/MAO.0000000000002313

Okayasu, T., Quesnel, A. M., O’Malley, J. T., Kamakura, T., & Nadol, J. B. (2020). The Distribution and Prevalence of Macrophages in the Cochlea Following Cochlear Implantation in the Human: An Immunohistochemical Study Using Anti-Iba1 Antibody. Otology and Neurotology, 41(3), e304–e316. https://doi.org/10.1097/MAO.0000000000002495

Parakalan, R., Jiang, B., Nimmi, B., Janani, M., Jayapal, M., Lu, J., Tay, S. S. W., Ling, E. A., & Dheen, S. T. (2012). Transcriptome analysis of amoeboid and ramified microglia isolated from the corpus callosum of rat brain. BMC Neuroscience, 13(1). https://doi.org/10.1186/1471-2202-13-64

Puntambekar, S. S., Saber, M., Lamb, B. T., & Kokiko-Cochran, O. N. (2018). Cellular players that shape evolving pathology and neurodegeneration following traumatic brain injury. In Brain, Behavior, and Immunity (Vol. 71, pp. 9–17). Academic Press Inc. https://doi.org/10.1016/j.bbi.2018.03.033

Quesnel, A. M., Nakajima, H. H., Rosowski, J. J., Hansen, M. R., Gantz, B. J., & Nadol, J. B. (2016). Delayed loss of hearing after hearing preservation cochlear implantation: Human temporal bone pathology and implications for etiology. Hearing Research, 333, 225–234. https://doi.org/10.1016/j.heares.2015.08.018

Rai, V., Wood, M. B., Feng, H., Schabla, N. M., Tu, S., & Zuo, J. (2020). The immune response after noise damage in the cochlea is characterized by a heterogeneous mix of adaptive and innate immune cells. Scientific Reports, 10(1), 1–17. https://doi.org/10.1038/s41598-020-72181-6

Roche, J. P., & Hansen, M. R. (2015). On the Horizon: Cochlear Implant Technology. Otolaryngologic Clinics of North America, 48(6), 1097–1116. https://doi.org/10.1016/j.otc.2015.07.009

Rowe, D., Chambers, S., Hampson, A., Eastwood, H., Campbell, L., & O’Leary, S. (2016). Delayed low frequency hearing loss caused by cochlear implantation interventions via the round window but not cochleostomy. Hearing Research, 333, 49–57. https://doi.org/10.1016/j.heares.2015.12.012

Sato, E., Shick, H. E., Ransohoff, R. M., & Hirose, K. (2008). Repopulation of cochlear macrophages in murine hematopoietic progenitor cell chimeras: The role of CX3CR1. Journal of Comparative Neurology, 506(6), 930–942. https://doi.org/10.1002/cne.21583

Sato, E., Shick, H. E., Ransohoff, R. M., & Hirose, K. (2010). Expression of fractalkine receptor CX3CR1 on cochlear macrophages influences survival of hair cells following ototoxic injury. JARO - Journal of the Association for Research in Otolaryngology, 11(2), 223–234. https://doi.org/10.1007/s10162-009-0198-3

Savage, J. C., Carrier, M., & Tremblay, M. È. (2019). Morphology of Microglia Across Contexts of Health and Disease. In Methods in Molecular Biology (Vol. 2034, pp. 13–26). Humana Press Inc. https://doi.org/10.1007/978-1-4939-9658-2_2

Scheperle, R. A., Tejani, V. D., Omtvedt, J. K., Brown, C. J., Abbas, P. J., Hansen, M. R., Gantz, B. J., Oleson, J. J., & Ozanne, M. V. (2017). Delayed changes in auditory status in cochlear implant users with preserved acoustic hearing. Hearing Research, 350, 45–57. https://doi.org/10.1016/j.heares.2017.04.005

Seyyedi, M., & Nadol, J. B. (2014). Intracochlear inflammatory response to cochlear implant electrodes in humans. Otology and Neurotology, 35(9), 1545–1551. https://doi.org/10.1097/MAO.0000000000000540

Shepherd, R. K., Carter, P. M., Enke, Y. L., Wise, A. K., & Fallon, J. B. (2019). Chronic intracochlear electrical stimulation at high charge densities results in platinum dissolution but not neural loss or functional changes in vivo. Journal of Neural Engineering, 16(2), aaf66b. https://doi.org/10.1088/1741-2552/aaf66b

Tan, W. J. T., Thorne, P. R., & Vlajkovic, S. M. (2016). Characterisation of cochlear inflammation in mice following acute and chronic noise exposure. Histochemistry and Cell Biology, 146(2), 219–230. https://doi.org/10.1007/s00418-016-1436-5

Wilk, M., Hessler, R., Mugridge, K., Jolly, C., Fehr, M., Lenarz, T., & Scheper, V. (2016). Impedance Changes and Fibrous Tissue Growth after Cochlear Implantation Are Correlated and Can Be Reduced Using a Dexamethasone Eluting Electrode. PLoS ONE, 11(2), 1–19. https://doi.org/10.1371/journal.pone.0147552

Wood, M. B., & Zuo, J. (2017). The contribution of immune infiltrates to ototoxicity and cochlear hair cell loss. Frontiers in Cellular Neuroscience, 11(April), 1–7. https://doi.org/10.3389/fncel.2017.00106

Wynn, T. A., & Vannella, K. M. (2016). Macrophages in Tissue Repair, Regeneration, and Fibrosis. Immunity, 44(3), 450–462. https://doi.org/10.1016/j.immuni.2016.02.015

